# PanCircBase: An online resource for the exploration of circular RNAs in pancreatic islets

**DOI:** 10.1101/2022.04.29.490033

**Authors:** Tanvi Sinha, Smruti Sambhav Mishra, Suman Singh, Amaresh Chandra Panda

**Affiliations:** Institute of Life Sciences, Nalco Square, Bhubaneswar, Odisha, India; Regional Center for Biotechnology, Faridabad, India

**Keywords:** Circular RNA, pancreatic islet, divergent primer, microRNA, translation

## Abstract

Circular RNAs (circRNAs) are a novel class of covalently closed RNA molecules that recently emerged as a critical regulator of gene expression in development and diseases. Recent research has highlighted the importance of novel circRNAs in the biosynthesis and secretion of insulin from β-cells of pancreatic islets. However, all circRNAs expressed in pancreatic islets or β-cells are not readily available in the database. In this study, we analyzed publicly available RNA-sequencing datasets of the pancreatic islets to catalog all circRNAs expressed in pancreatic islets to construct the PanCircBase (www.pancircbase.net/) database that provides the following resources: (i) pancreatic islet circRNA annotation details (genomic position, host gene, exon information, splice length, sequence, other database IDs, cross-species conservation), (ii) divergent primers for PCR analysis of circRNAs, (iii) siRNAs for silencing of target circRNAs, (iv) miRNAs associated with circRNAs, (v) possible protein-coding circRNAs and their polypeptides. In summary, this is a comprehensive online resource for exploring circRNA expression and its possible function in pancreatic β-cells.

## 1. Introduction

The last few decades have seen a significant increase in the incidences and prevalence of diabetes around the globe [1]. Diabetes is often associated with a decline in insulin sensitivity or pancreatic β-cell dysfunction with decreased insulin production to maintain glucose homeostasis [2]. In normal conditions, pancreatic β-cells increase insulin production in response to increased glucose to maintain glucose homeostasis. However, prolonged exposure of pancreatic β-cells with high glucose or high fat leads to a decrease in insulin production by β-cell and causes diabetes [2,3]. Several studies established that the β-cell gene expressions are regulated at transcriptional and posttranscriptional levels. Interestingly, the genome-wide association studies identified more than a hundred type2 diabetes-associated risk loci and the majority of them reside in the noncoding part of the genome [4]. Although most of the human genome is pervasively transcribed into RNA, only a fraction of it translates into proteins [5,6]. The noncoding RNA family contains several types of RNAs and is mainly involved in regulating protein-coding genes. Recent studies established that long noncoding RNAs and microRNAs (miRNAs) regulate normal β-cell function and are involved in the development of diabetes [7-9]. However, the role of recently discovered circular RNA (circRNA) molecules in pancreatic β-cell function has not been understood completely [7,10].

Discovered more than forty years ago, circRNAs were believed to be artifacts or splicing errors generating head-to-tail covalently closed RNAs without significant physiological importance [11-13]. In the last decade, the dramatic improvements in novel computational pipelines and high throughput transcriptomic sequencing technology made it possible to realize the universal expression and function of circRNAs [14-16]. In contrast to messenger RNAs (mRNAs), circRNAs are a novel class of ubiquitously expressed closed-loop single-stranded RNA molecules without the cap and poly-A tail. Since circRNAs are highly stable, they can act as sponges for miRNAs and RNA-binding proteins (RBPs) [17,18]. Although circRNAs are mostly categorized as noncoding RNAs, a few have been reported to translate into proteins [19]. In addition, the secretion of stable circRNAs into the body fluid or exosomes makes them a promising biomarker for disease diagnosis [20]. Hundreds of recent studies suggest that circRNAs are associated with disease development and progression, including cancer, cardiovascular, rheumatoid arthritis, Alzheimer’s, and diabetes [21]. Recently, a few studies highlighted the role of circRNAs in pancreatic β-cell physiology and the development of diabetes [22-25]. However, the complete landscape of circRNA expression in pancreatic islets and their involvement in diabetes is still not well understood.

Given the emerging physiological importance of circRNAs and research efforts to understand them in pancreatic islet physiology, we developed the circRNA repository called PanCircBase to catalog the circRNAs expressed in pancreatic islets and provide resources for further research (Figure 1). This database catalogs known and novel circRNAs expressed in pancreatic islets. Given that circRNAs regulate cellular physiology, PanCircBase provides precise information about divergent primers, siRNAs, and miRNA targets for functional analysis of pancreatic islet circRNAs. In addition, the database provides information on the possible translation of circRNAs into proteins. Together, PanCircBase provides interactive tools to access pancreatic islet circRNA expression as well as information for functional validation of circRNAs to understand their role in ß-cell function and diabetes.

**Figure 1.**
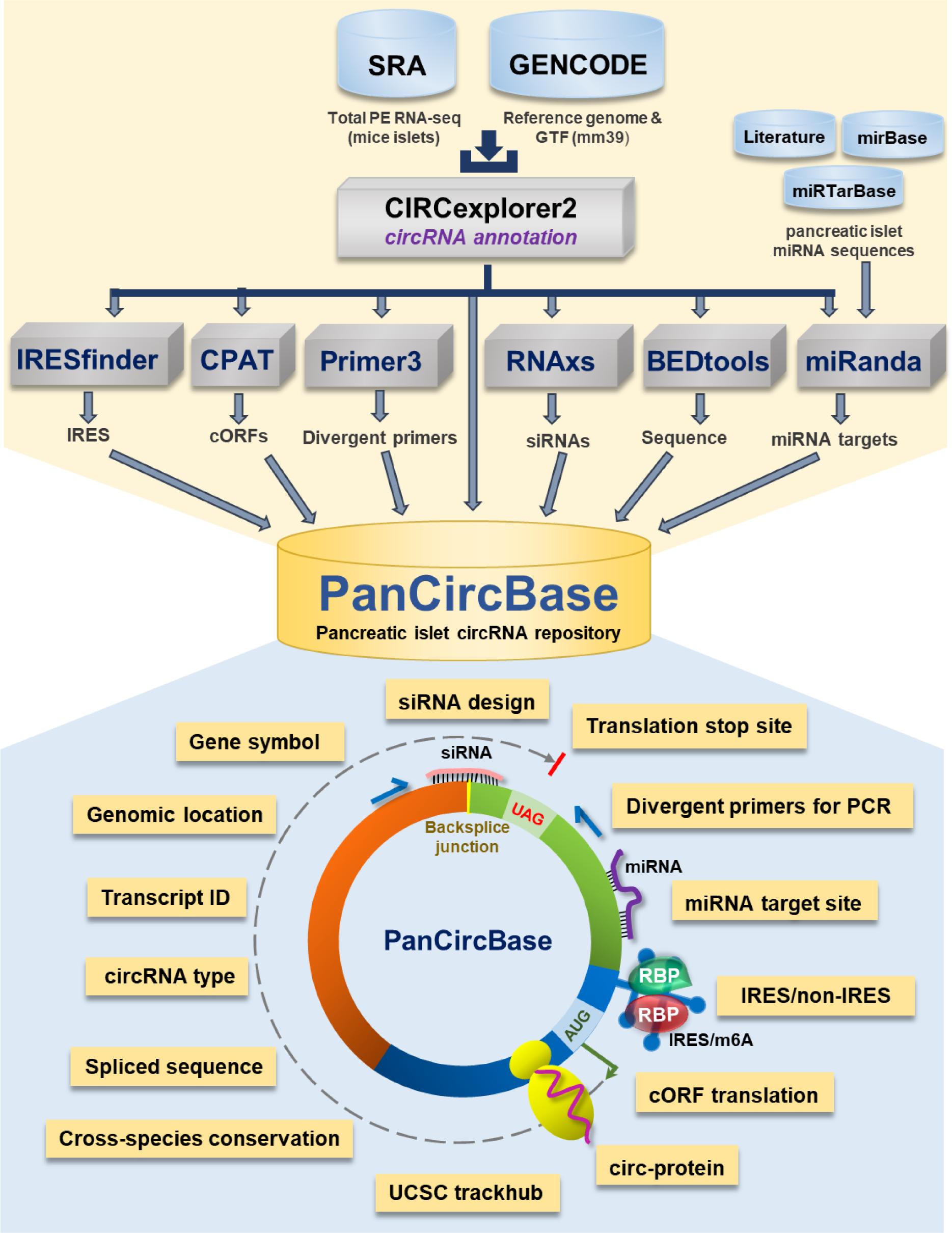
The framework of pancreatic islet circRNA database “PanCircBase”.

## 2. Results

### 2.1. Pancreatic islet circRNAs and their properties

The publicly available total RNA-seq data from mice pancreatic islets were analyzed with CIRCexplorer2 pipeline to identify circRNAs expressed in pancreatic islets (NCBI-SRA BioProject Accession No: PRJNA681104 and PRJNA358100) [26-28]. We identified 66,501 circRNAs in pancreatic islets, out of which 90% were shorter than 2000 nucleotides in length (Figure 1A, Supplemental Table S1). Identification of circRNA host genes revealed that nearly 16% of genes expressed in pancreatic islets also produce circular RNAs (Figure 2B). Since circRNAs are produced from intronic and exonic sequences, pancreatic islet circRNA annotation suggested that only 8% of the circular RNAs are generated solely from the intronic regions while 92% circRNAs were generated from exons (Figure 2C). The complete list of the 66,501 circRNAs along with their chromosomal coordinates, circRNA host gene information, length, and normalized expression of circRNAs in normal mice islets or high-fat-fed mice or db/db mice are provided in Supplemental Table S1. In addition, we identified the islet circRNAs in other databases and mentioned their IDs from circBase, CIRCpedia v2, and circAtlas v2.0 in the web interface [29-31]. Interestingly, more than 23,000 novel circRNAs identified in pancreatic islets were novel circRNAs, not reported in the above databases. Furthermore, we performed the conservation analysis of mouse pancreatic islet circRNAs by converting the genomic coordinates of circRNAs to the human genome coordinates using the UCSC liftOver tool and analyzed their expression in human samples in circBase, CIRCpedia v2, and circAtlas v2.0 databases [29-32]. Notably, more than half of the pancreatic islet circRNAs were reported to be expressed in human samples in the above databases suggesting their conservation in humans.

**Figure 2:**
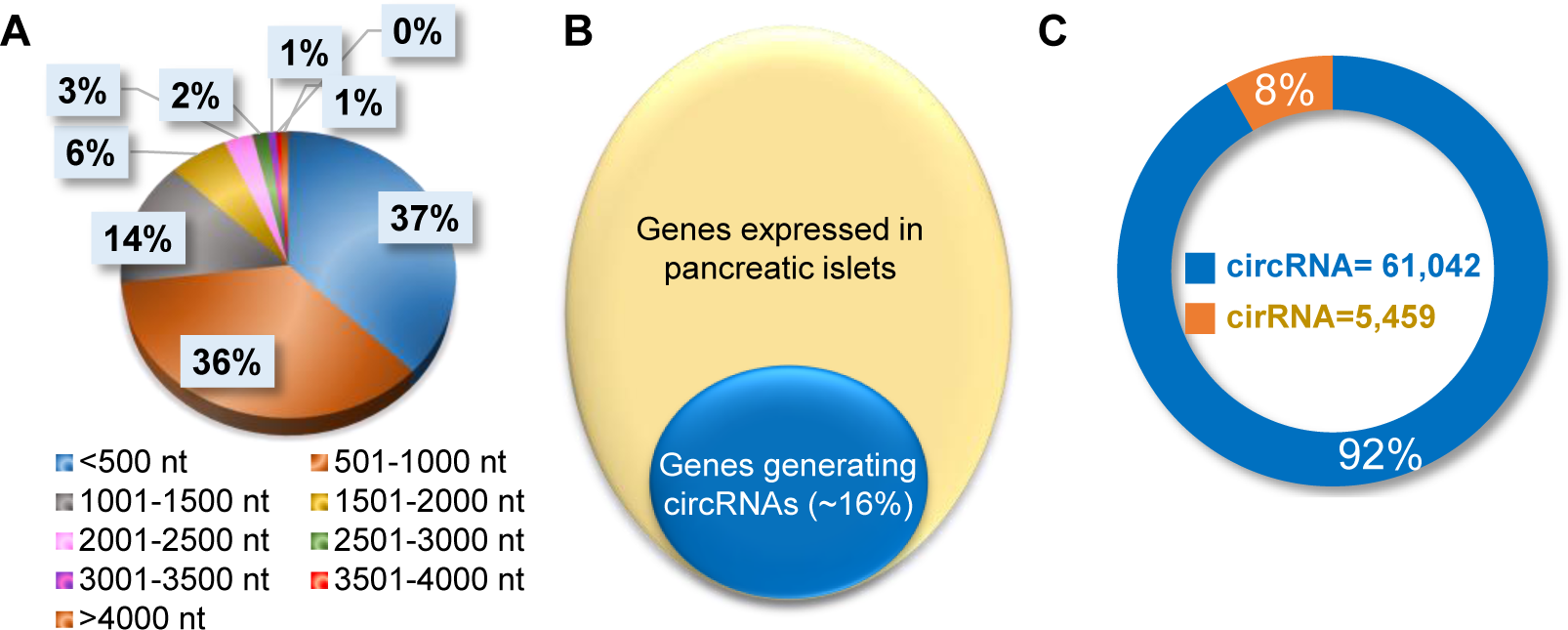
Characteristics of circRNAs expressed in pancreatic islets. **A**.. Percentage distribution of length of circRNAs in pancreatic islets. **B**. Percentage of genes generating circRNAs in pancreatic islets. **C**. Number of circular RNAs categorized as exonic circRNA or intronic ciRNAs in pancreatic islets.

### 2.2. PanCircBase search function & detailed information on each circRNAs

Here, the circRNAs are named based on the host gene and the mature length of the circRNAs (PanCircBase ID: gene name_circRNA length in nt). For example, the PanCircBase ID for the exonic circular (circ)RNA of 243 nt length from the *Ankrd12* gene is “*mmu_circAnkrd12_243*”. The web interface of PanCircBase is shown in Figure 3. The home page of PanCircBase contains the introduction message, related publications, and a “quick search window” for searching circRNAs by their host gene name. If a gene symbol is queried, the search results enlist all the circRNAs originating from the gene of interest in pancreatic islets. For example, if the user wants to search for circRNAs generated from the *Ankrd12* gene, upon inputting the keyword “*Ankrd12*”, PanCircBase search results show all circRNAs originating from the *Ankrd12* gene such as *mmu_circAnkrd12_243, mmu_circAnkrd12_286, circAnkrd12_339, circAnkrd12_344*, etc. The user can get a basic annotation of each circRNA such as PanCircBase ID, chromosome position, host gene name, transcript ID, genomic length, mature spliced length, and circRNA sequence by clicking the “Quick view”. The user can click the “Full details” to find the detailed information on circRNA, including PanCircBase ID, annotation details, divergent primers, miRNAs, siRNAs, and protein-coding potential details of circRNA (Figure 3).

**Figure 3.**
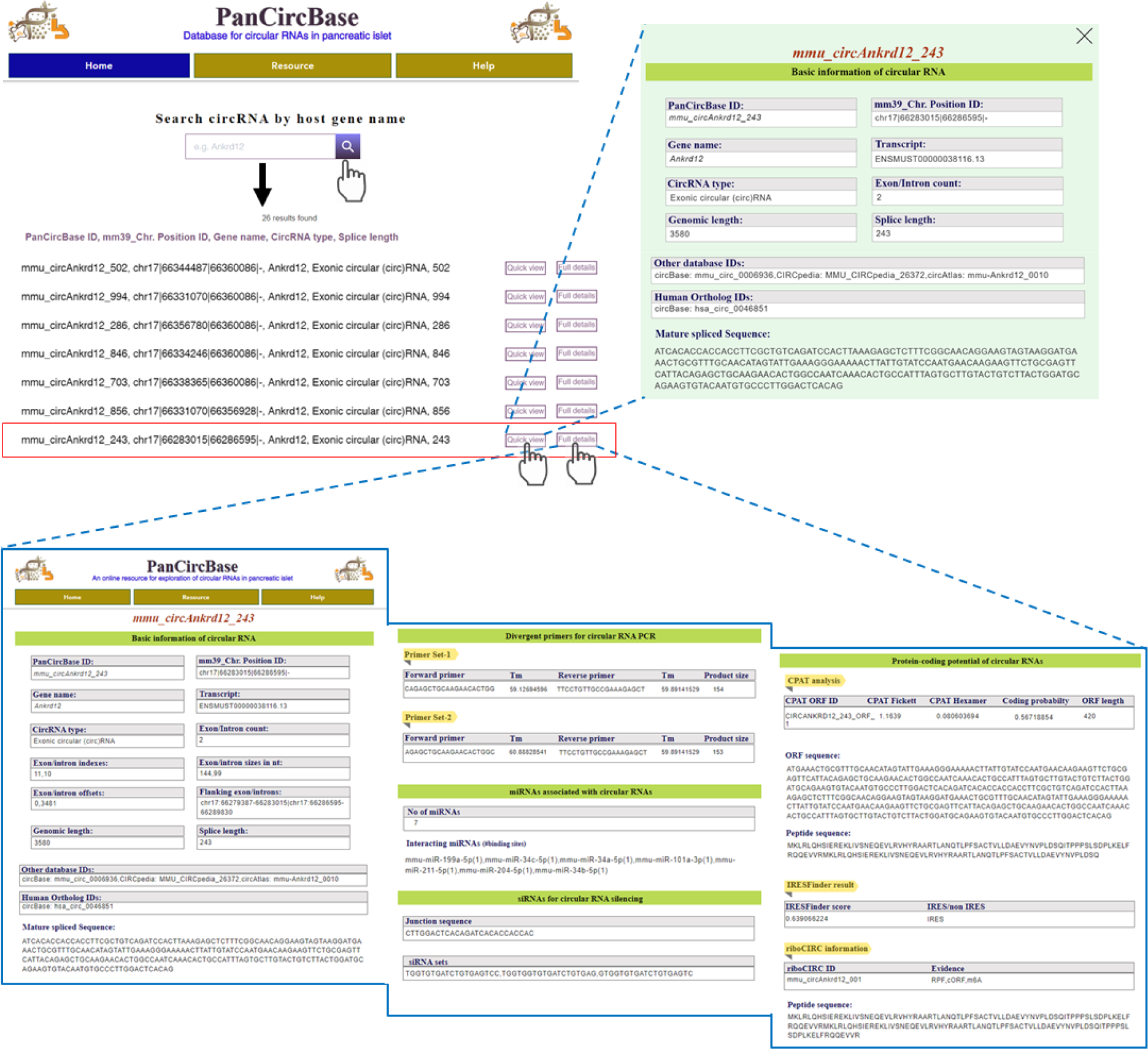
Overview of PanCircBase web interface. **A**.. Users can search by the host gene symbol “Ankrd12,” The search results show all the circRNAs originated from the *Ankrd12* gene. When users click “Quick view”, PanCircBase opens a new window with the basic information on selected circRNA, and clicking the “Full details” opens a new page with details of selected circRNA, including circRNA annotation, PCR primers, siRNAs, target miRNAs, and possible protein-coding ability of the circRNAs.

### 2.3. Resources for circRNA functional analysis

PanCircBase is a user-friendly and comprehensive database that provides useful information on 66,501 circRNAs expressed in mouse pancreatic islet. Besides the detailed circRNA information, PanCircBase also provides other information for the functional characterization of circRNAs, including divergent primers, siRNAs, miRNAs, and polypeptides encoded by the pancreatic islet circRNA.

#### 2.3.1. Divergent primer for circRNA PCR

A few web tools are available for designing primers for circRNAs present in their database. For example, circInteractome can design divergent primers only for human circRNAs listed in circBase [33]. However, currently no software or web server is available for designing divergent primers for novel circRNAs. Here, we provide two sets of specific divergent primer pairs for validation and quantification of circRNA using reverse transcription (RT) followed by PCR (RT-PCR) analysis (Supplementary Table S2). The divergent primer pairs are designed for specific PCR amplification of the target circRNA backsplice junction sequence, which can be further validated by Sanger sequencing using one of the divergent primers (Figure 3).

#### 2.3.2. siRNA for circRNA silencing

The lack of appropriate siRNA designing tools and the unavailability of circRNA junction sequences make it challenging to design siRNAs against circRNAs. Thus, we designed siRNAs targeting each pancreatic islet circRNAs using the RNAxs siRNA design tool [34]. PanCircBase provides 19-nt siRNAs targeting backsplice junction sequence spanning a minimum of six nucleotides on either side of the junction. We could successfully design 1 to 3 siRNAs for 43,483 circRNAs in the PanCircBase (Supplementary Table S3). For example, three siRNA sequences targeting the backsplice junction sequence were predicted for *mmu_circAnkrd12_243* (Figure 3). The user needs to confirm the specificity of the siRNA using NCBI-BLAST. The siRNA sequences may be custom synthesized with two additional nucleotides (dTdT) as 3’ DNA overhangs.

#### 2.3.3. Predicting functional miRNA targets

Evidence suggests that circRNAs regulate gene expression by binding to miRNAs and RBPs. At present, circRNAs regulating gene expression through the circRNA-miRNA-mRNA regulatory network is the most extensively studied and accepted mechanism of circRNA-mediated gene regulation. Here, we downloaded functional miRNAs from miRTarBase and searched for pancreatic islet miRNA expression data in the literature [35-40]. We retrieved the mature sequences of 478 functional miRTarBase miRNAs expressed in pancreatic islets from miRBase (Supplementary Table S4) [41]. Systematic miRNA target site search on the pancreatic islet circRNAs using the miRanda program identified 63,505 circRNAs with at least one miRNA binding site [42]. For example, *mmu_circAnkrd12_243* harbors binding sites for seven miRNAs, and each miRNA is predicted to have one binding site on *mmu_circAnkrd12_243* (Figure 3). Furthermore, we discovered that 215 circRNAs with more than 5 binding sites for a single miRNA and may act as miRNA sponges (Supplementary Table S4).

#### 2.3.3. Protein-coding potential of islet circRNAs

Although circRNAs do not contain 5’ cap and poly-A tail for conventional translation, many recent reports suggested the association of hundreds of circRNAs with translation polyribosomes [19]. Furthermore, ribosome footprinting assays also identified circRNAs associated with ribosomes, suggesting possible translation of circRNAs into protein products through cap-independent mechanisms [19]. Several studies have established that m6A marks or internal ribosome entry sites (IRES) on circRNA sequences promote cap-independent translation of circRNAs into functional polypeptides [43-45]. The Coding-Potential Assessment Tool (CPAT) analysis identified 59,580 circRNAs with at least 1 ORF spanning the backsplice junction, suggesting that islet circRNAs could translate into proteins (Supplementary Table S5) [46]. We could identify 40,146 pancreatic islet circRNAs with potential IRES based on IRESfinder analysis (Supplementary Table S6) [47]. For example, *mmu_circAnkrd12_243* is predicted to have IRES by IRESfinder. In addition, we found that 475 circRNAs in PanCircBase can translate into proteins based on riboCIRC data (Supplementary Table S7) [48]. Interestingly, the riboCIRC database suggests that *mmu_circAnkrd12_243* (riboCIRC ID: *mmu_circAnkrd12_001)* can be translated into proteins based on pieces of evidence such as RPF, cORF, and m6A. Since we used a 2x circRNA sequence, the peptide sequence is truncated at the end of the circRNA sequence, while riboCIRC uses 4x circRNA sequence, giving a longer peptide sequence.

#### 2.3.4. UCSC track for circRNAs visualization

For direct visualization of pancreatic islet circRNAs in the UCSC genome browser, we created a link for the UCSC track of the circRNAs expressed in pancreatic islets. The user can use the link for the UCSC track provided on the resource page of PanCircBase to visualize the circRNAs generated from the gene of interest in pancreatic islets and circRNAs reported by circBase. For example, circBase reported the expression of five circRNAs from the *Ankrd12* gene, while PanCircBase identified twenty-six circRNAs generated from the *Ankrd12* gene, including *mmu_circAnkrd12_243* (circBase ID: mmu_circ_0006936) (Figure 4).

**Figure 4.**
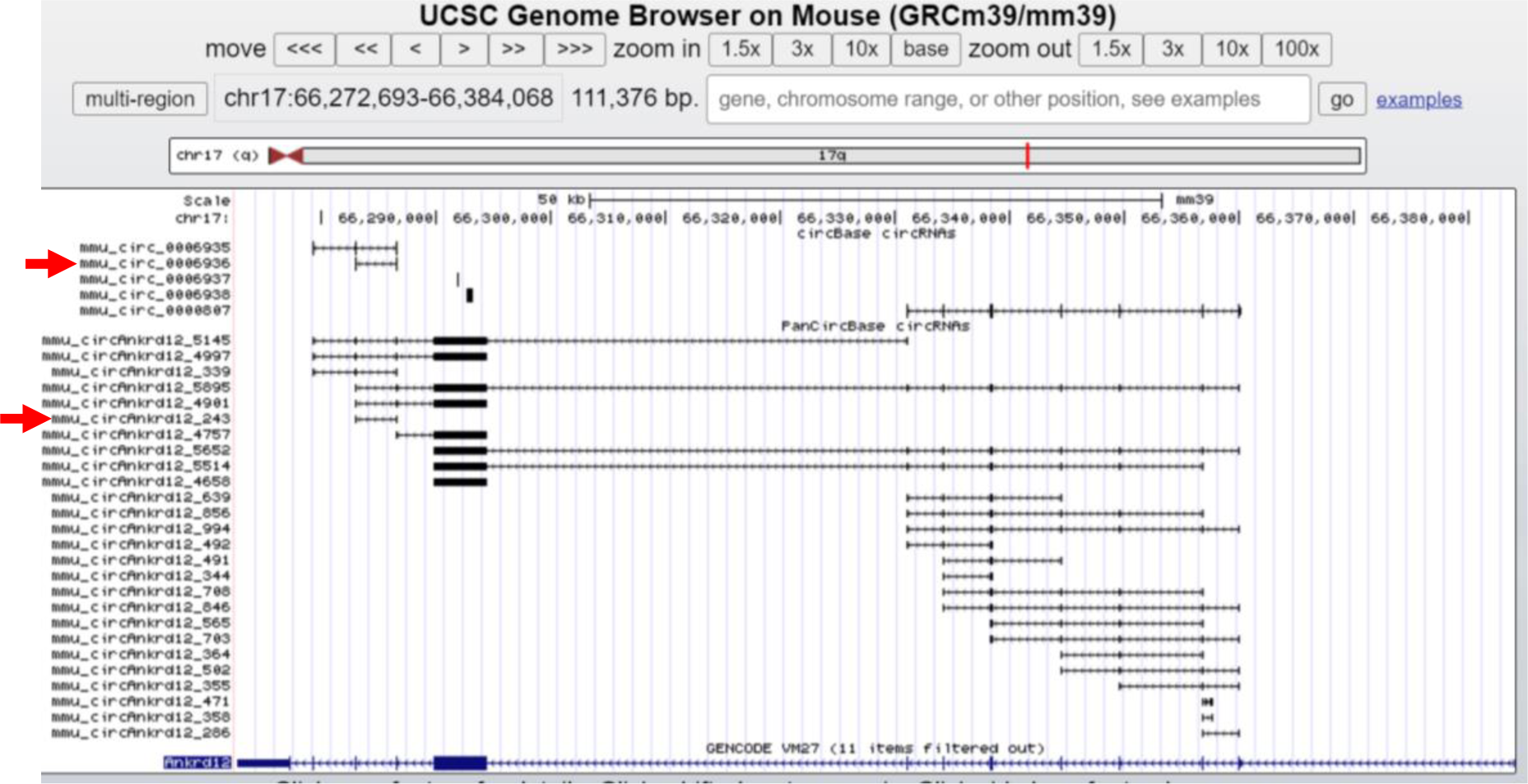
Overview of circRNAs generated from Ankrd12 in UCSC genome browser. The red arrow indicates the *mmu_circAnkrd12_243* and the corresponding circBase ID.

## 3. Discussion

The PanCircBase is a user-friendly and comprehensive database that can facilitate the study of circRNA expression and its function in pancreatic islets. PanCircBase comprises collection of publicly available RNA-seq data along with information from several computational tools and databases for circRNA analysis, including CIRCexplorer2, Primer3, RNAxs, miRTarBase, miRBase, miRanda, IRESfinder, CPAT, circBase, circAtlas, CIRCpedia, and riboCIRC. PanCircBase provides users with pancreatic islet circRNA expression levels, mature spliced sequences, circRNA-associated miRNAs, siRNAs, and divergent primers to analyze circRNAs. PanCircBase also provides two sets of divergent primer pairs for PCR validation and quantification of circRNA. Since circRNA silencing is one of the most relevant ways to study their function in physiological conditions, PanCircBase provides a few siRNA sequences targeting the backsplice junction sequence for specific circRNA silencing. Furthermore, the presence of open reading frames along with IRES on pancreatic islet circRNAs suggests their translatability into functional proteins.

Although PanCircBase provides a user-friendly platform for pancreatic islet circRNA research, it also enlists the circRNAs expressed in the whole islet that includes insulin-producing β-cells along with other non-insulin-producing cells. Therefore, some of the circRNA reported here could be expressed by non-insulin-producing cells as well. Furthermore, since the RNA-sequencing was for total RNA without depletion of linear RNAs and the circRNA sequences were predicted based on the mm39 genome annotation, the expression of circRNA splice variants with different exon/intron combinations cannot be ruled out. Given that the circRNA-associated miRNAs were predicted based on the miRanda program computationally, biochemical experiments are essential to verify functional circRNA-miRNA interaction and their downstream target gene regulation.

We will continue to maintain and update the features of PanCircBase in the foreseeable future as the new data become available. We will include circRNAs expressed in pure β-cells or in different physiological conditions as the more relevant data becomes available. Furthermore, we plan to include experimentally validated functional pancreatic islet circRNAs and their regulatory network in PanCircBase. We also plan to integrate pancreatic islet or β-cell circRNAs from other species such as humans and will also include the circRNA-miRNA/RBP regulatory networks. Together, PanCircBase is a valuable resource to accelerate understanding of the physiological relevance of pancreatic islet circRNAs in diabetes.

## 4. Materials and Methods

### 4.1. RNA-seq data collection and circRNA analysis

We collected published total RNA-seq data of pancreatic islets of normal mice, db/db mice, or mice fed with a normal and high-fat diet (NCBI-SRA BioProject Accession No: PRJNA681104 and PRJNA358100) [26-28]. The quality of the RNA-seq reads was checked with FastQC software (v0.11.9). STAR aligner (v2.7.10a) was used to align the RNA-seq reads to the mm39 mouse genome using the ChimSegmentMin −10 parameter. The chimeric reads from the STAR aligner were used to identify the circRNAs in pancreatic islets using the CIRCexplorer2 pipeline (v2.3.8) [28]. The CIRCexplorer2 pipeline provides basic genomic information of the circRNA in BED format, including chromosome number, chromosome start, chromosome end, strand, host gene symbol, transcript ID, exon information, and exon length. Furthermore, we retrieved the mature spliced circRNA sequences using BEDtools (v2.29.1). Since the pancreatic islet RNA-seq data were collected from different sources and the sequencing depth was different for different samples, we represent normalized circRNA expression levels as transcripts per million (TPM: circRNA read number/total circRNA reads in the sample x 1,000,000) along with the raw read numbers from one set of sample. The circRNA annotation data and the expression levels are provided in Supplementary Table S1. In addition, we also retrieved the IDs for pancreatic islet circRNAs in various databases such as circBase, circAtlas v2.0, CIRCpedia v2, and riboCIRC v1.0 [29-31,48]. In addition, the conservation of PanCircBase circRNAs was analyzed by converting the circRNA chromosomal locations to human coordinates using the UCSC liftOver [32]. The converted human coordinates were searched in circBase, circAtlas v2.0, and CIRCpedia v2 databases to find the human orthologs of PanCircBase circRNAs.

### 4.2. Divergent primer and siRNA design

The PCR amplicon template of circRNA backsplice junction sequences was prepared by joining 100 nt from either side of the mature sequence as described in our previous publication [49]. The PCR amplicon was prepared for circRNAs shorter than 200 nt by joining the 3’ half to the 5’ half sequence. We used the Python package for the Primer3 oligo design tool (primer3-py) for designing divergent primer pairs that can amplify the backsplice junction with a product ranging from 120-180 bp for circRNAs longer than 120 nt, and PCR products of 60-120 bp for circRNAs less than 120 nt length [50]. Also, we used the Tm cutoff range 55^0^ C-63^0^ C and the primer length range 18-27 nt. CircRNAs with less than 60 nt lengths were not taken for primer design.

The backsplice junction sequence of 26 nt spanning 13 nt on either side of the circRNA junction was derived from the mature spliced sequence. The 26 nt junction sequence was used by the RNAxs tool as input for designing 19-nt siRNAs against circRNA junction sites with a minimum of 6 nt on either side of the backsplice junction [34].

### 4.3. Prediction of associated functional miRNAs

The miRNAs expressed in mouse pancreatic islets or β-cells were derived from previous publications, and the validated mouse functional miRNAs were retrieved from the miRTarBase release 8 (Supplementary Table S4) [35]. The miRNAs with validated target genes in the miRTarBase and expressed in mouse pancreatic islets were selected for studying their association with pancreatic islet circRNAs [35]. The mature miRNA sequences were downloaded from the miRBase database [41]. The mature circRNAs sequences were used in miRanda v3.3a software to predict the functional islet miRNAs associated with target circRNAs [42].

### 4.4. Potential protein-coding circRNAs

Here, we used the CPAT v3.0.4 to analyze the protein-coding potential of pancreatic islet circRNAs [46]. The CPAT scores positively correlate to the protein-coding potential of the circRNA (Supplementary Table S5). Since circRNAs are known to be translated through IRES-mediated translation initiation, we also used the IRESfinder v1.1.0 tool to predict the IRES element of pancreatic islet circRNAs (Supplementary Table S6) [47]. In addition, the list of protein-coding mouse circRNAs was downloaded from the riboCIRC database [48]. The pancreatic islet circRNAs with protein-coding ability were identified from the riboCIRC database, their potential peptide sequence, and the presence of IRES/m6A/mass spectrometry data as supporting evidence (Supplementary Table S7).

## Supplementary Materials

**Supplementary Table S1**. Annotation details of circRNAs expressed in pancreatic islets. **Supplementary Table S2**. Divergent primer pairs for PCR amplification of circRNAs. **Supplementary Table S3**. List of siRNAs for silencing of circRNAs expressed in pancreatic islets. **Supplementary Table S4**. MicroRNAs targeting circRNAs expressed in pancreatic islets. **Supplementary Table S5**. The protein-coding potential of circRNAs predicted by CPAT. **Supplementary Table S6**. Prediction of IRES-containing circRNAs using IRESfinder. **Supplementary Table S7**. List of protein-coding islet circRNAs in riboCIRC database.

## Author Contributions

Conceptualization, T.S., S.S.M., and A.C.P; methodology, T.S. and S.S.M.; software, T.S., and S.S.M.; validation, T.S., S.S., and S.S.M.; formal analysis, T.S., S.S., and S.S.M.; investigation, T.S. and S.S.M.; writing—original draft preparation, T.S., and A.C.P; writing—review and editing, T.S., S.S.M.,S.S., and A.C.P; visualization, T.S. and A.C.P.; supervision, A.C.P.; funding acquisition, A.C.P. All authors have read and agreed to the published version of the manuscript.

## Acknowledgments

The authors thank Arup Ghosh and Sharmishtha Shyamal for helpful discussions and critical proofreading of this manuscript.

## Data Availability Statement

The PanCircBase is now publicly accessible at www.pancircbase.net/. All data generated in this study are included in the main text, database, or supplementary information files. The circRNAs identified in this study with their detailed annotations, expression levels, and mature spliced sequences can be directly downloaded from the database. In addition, the circRNA-miRNA interaction data with the scores can be downloaded from the download page of the database.

